# ProtAttn-QuadNet: An attention-based deep learning framework for protein–protein interaction prediction using ProtBERT embeddings

**DOI:** 10.1101/2025.11.17.688786

**Authors:** Md. Shahidul Islam, Md. Muhtasim Rahman Mim, Md. Raihan Kabir

**Affiliations:** Department of Computer Science and Engineering, University of Asia Pacific, 74/A, Greenroad, Dhaka-1205, Bangladesh

## Abstract

Protein–protein interactions (PPIs) form the backbone of most cellular processes, governing signal transduction, gene regulation, and metabolic control. However, experimental approaches to identifying PPIs remain expensive, laborious, and often incomplete. Recent advances in protein language models (PLMs) have transformed sequence-based PPI prediction by enabling deep contextual encoding of biochemical and structural information directly from amino acid sequences. Building upon this progress, we present ProtAttn-QuadNet, an attention-based deep learning framework that leverages ProtBERT embeddings to model reciprocal dependencies between protein pairs. The proposed model employs a quad-stream attention mechanism that integrates individual protein features, synergistic interactions, and complementary differences through multi-level self- and cross-attention layers. This architecture enables the discovery of fine-grained relational patterns while ensuring balanced bidirectional modeling of interacting proteins. Evaluated on large-scale dataset from UniProt, ProtAttn-QuadNet achieves 97.16% accuracy (AUC-ROC 99.00%) on balanced data and 99.19% accuracy (AUC-ROC 99.76%) on oversampled datasets, surpassing several recent state-of-the-art PPI prediction methods. Statistical validation using the Chi-square and Wilcoxon signed-rank tests confirms the model’s predictive significance and reliability. ProtAttn-QuadNet offers a powerful computational framework for large-scale PPI prediction.

**Author summary:** Proteins work together to carry out almost every function in living cells, from sending signals to controlling metabolism. Knowing which proteins interact with each other helps scientists understand how cells work and how diseases develop. However, finding these interactions in the laboratory is often slow, costly, and incomplete. In this study, a computational model called ProtAttn-QuadNet is developed to predict protein–protein interactions using only the amino acid sequences of proteins. The model analyses each pair of proteins to find shared features and differences that indicate whether they interact. It also uses a set of attention layers that allow the model to focus on the most relevant sequence patterns. When tested on a large protein dataset, ProtAttn-QuadNet produced highly accurate and consistent results, performing better than several existing methods. These results suggest that ProtAttn-QuadNet can serve as a reliable tool for studying protein networks and may help guide future research in biology, medicine, and drug development.

## Introduction

Protein–protein interactions (PPIs) are fundamental to almost all cellular processes, including signal transduction, gene expression regulation, metabolic control, and immune responses [1–3]. Understanding the complex network of PPIs provides valuable insights into cellular functions and disease mechanisms [4, 5]. Although numerous experimental techniques, such as yeast two-hybrid screening, co-immunoprecipitation, and affinity purification coupled with mass spectrometry, have been developed to detect PPIs, these methods remain time-consuming, costly, and often limited in coverage [6]. Consequently, computational prediction methods have become indispensable for large-scale PPI analysis.

Early computational approaches primarily relied on handcrafted sequence features, including amino acid composition, evolutionary profiles, and physicochemical descriptors. Classical machine learning algorithms such as Support Vector Machines (SVM), Random Forests (RF), and Bayesian classifiers were employed to classify interacting protein pairs based on these features [7–12]. While these models demonstrated moderate success, their dependence on manually engineered descriptors and incomplete structural data limited their generalization capabilities, particularly across species and diverse protein families [11, 13–15].

The increasing availability of large-scale protein sequence databases has encouraged sequence-based prediction methods that rely less on structural information. Deep learning has substantially advanced this field by enabling hierarchical feature extraction and representation learning. DeepPPI [16] used a fully connected neural network to model complex non-linear relationships between protein features, whereas DPPI [17] applied a Siamese-like convolutional architecture to learn symmetric relationships between interacting proteins. Similarly, PIPR [18] introduced a residual recurrent convolutional neural network (RCNN) to capture both local motifs and long-range dependencies, while Wu et al. proposed DL-PPI [19], a graph neural network–based model that integrates multi-scale features and attention mechanisms to enhance relational reasoning among proteins. These architectures collectively improved predictive performance but often struggled with interpretability, data imbalance, and computational efficiency.

Recent advances in transformer architectures and PLMs have transformed sequence-based PPI prediction by learning contextualized residue representations through self-attention mechanisms. Pretrained models such as ProtTrans [5], ProtBERT [20], and ESM-2 [21] encode rich biochemical and evolutionary information from massive unlabeled protein corpora, effectively capturing secondary and tertiary structure tendencies directly from primary sequences. Several recent studies have leveraged these embeddings for PPI prediction using hybrid deep architectures. For example, xCAPT5 [22] integrated ProtTrans embeddings with a multi-kernel convolutional network to capture local and global dependencies, while TUnA [23] incorporated uncertainty modeling within a transformer framework to improve robustness. PPI-Graphomer [24] combined pretrained language models with graph transformers to integrate sequence and structural representations, achieving high performance across benchmark datasets.

Despite these advances, existing frameworks often treat protein pairs asymmetrically and fail to explicitly model the reciprocal dependencies inherent in protein–protein binding. Moreover, many attention-based methods focus on single-sequence encoding and overlook bidirectional relationships critical to interaction dynamics.

To address these limitations, this study proposes ProtAttn-QuadNet, an attention-based deep learning framework that leverages ProtBERT embeddings to model mutual interactions between protein pairs. By incorporating quadratic attention mechanisms and a reciprocal representation module, ProtAttn-QuadNet identifies key sequence regions contributing to binding affinity and ensures balanced interaction assessment. The proposed model was evaluated on dataset collected from UniProt [25] and demonstrated high predictive accuracy and robustness across species. Overall, ProtAttn-QuadNet offers a reliable computational framework for large-scale PPI prediction and contributes to a deeper understanding of cellular interaction networks.

## Results and Discussions

### Evaluation Metrics

Evaluating the performance of our protein–protein interaction (PPI) prediction model is essential for assessing its ability to identify true interactions, minimize false predictions, and generalize to unseen protein pairs. We employed five standard performance metrics: accuracy, precision, recall, F1-score, and AUC-ROC. These metrics collectively assess the model’s overall correctness, reliability of positive predictions, sensitivity to true interactions, and balance between precision and recall. Performance was monitored across all training epochs to ensure stable learning and robust generalization.

- **Accuracy:** Measures the overall proportion of correctly predicted interacting and non-interacting protein pairs:

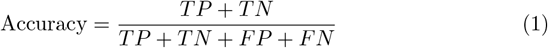
- **Precision:** Represents the fraction of correctly predicted interacting pairs among all predicted as interacting:

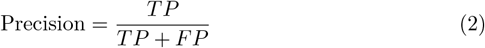
- **Recall (Sensitivity):** Quantifies the proportion of true interacting pairs correctly identified by the model:

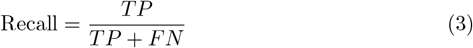
- **F1-Score:** The harmonic mean of precision and recall, providing a balanced measure of predictive performance:

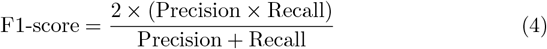
- **AUC-ROC:** Evaluates the trade-off between true positive and false positive rates across thresholds, reflecting the model’s ability to distinguish interacting from non-interacting pairs.

### Performance Evaluation of the Proposed Model

We evaluated the performance of the proposed protein–protein interaction (PPI) prediction model using two different datasets: a balanced dataset and an oversampled dataset. The balanced dataset allows us to assess how well the model performs when interacting and non-interacting protein pairs are equally represented, while the oversampled dataset increases the number of rare interactions, enabling us to evaluate how the model handles imbalanced data scenarios.

The comparative summary of model performance is presented in Table 1, highlighting the improvement gained through oversampling and confirming the proposed model’s reliability for large-scale PPI prediction tasks.

**Table 1.**
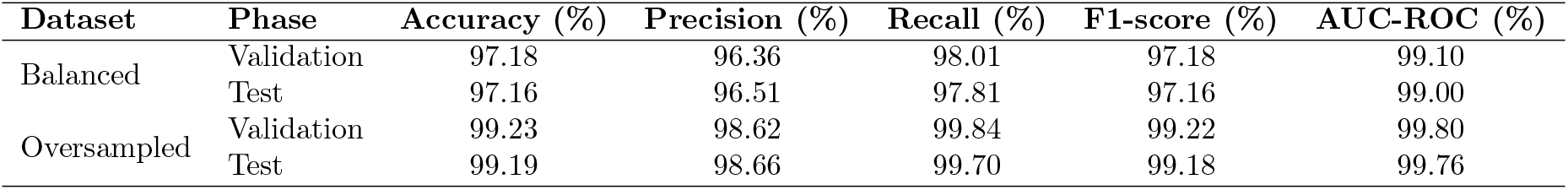
Comparison of model performance on balanced and oversampled datasets.

On the balanced dataset, the model achieved a validation accuracy of 97.18% and a test accuracy of 97.16%. The corresponding precision, recall, F1-score, and AUC-ROC were 96.36%, 98.01%, 97.18%, and 99.10% for validation, and 96.51%, 97.81%, 97.16%, and 99.0% for the test set, respectively. These metrics indicate that the model can effectively identify true interactions while maintaining a low false-positive rate. The high AUC-ROC demonstrates strong discriminative power, confirming that the model accurately differentiates between interacting and non-interacting protein pairs. The validation curves (Fig 1) further show consistent and stable learning, suggesting minimal overfitting and good generalization to unseen data.

**Fig 1.**
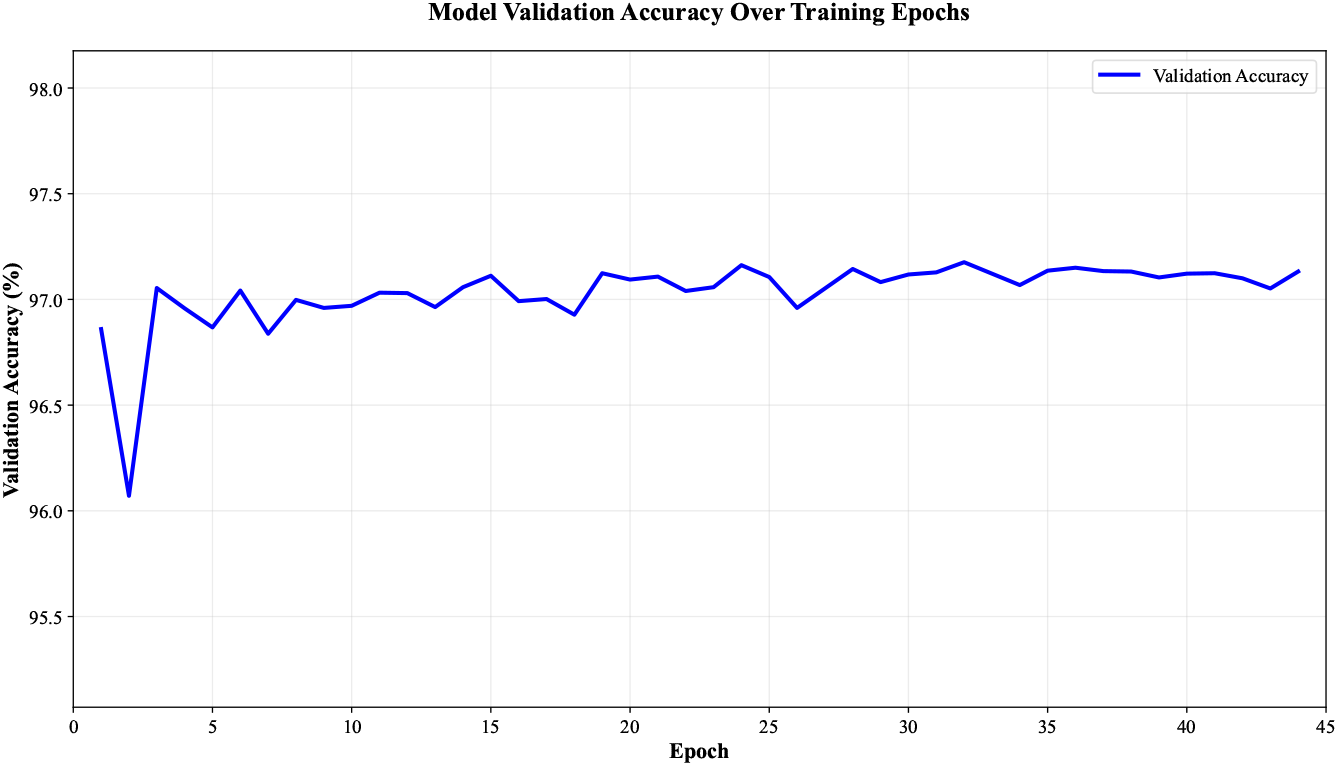
Validation Accuracy curve during training on the balanced dataset.

When trained on the oversampled dataset, the model demonstrated even higher performance. The validation accuracy reached 99.23% and the test accuracy 99.19%, while precision, recall, F1-score, and AUC-ROC reached 98.62%, 99.84%, 99.22%, and 99.80% for the validation set, and 98.66%, 99.70%, 99.18%, and 99.76% for the test set, respectively. The accuracy progression shown in Fig 2 illustrate a stable improvement throughout training. These findings suggest that the oversampling strategy effectively enhanced the model’s capacity to learn from minority interaction samples without introducing overfitting.

**Fig 2.**
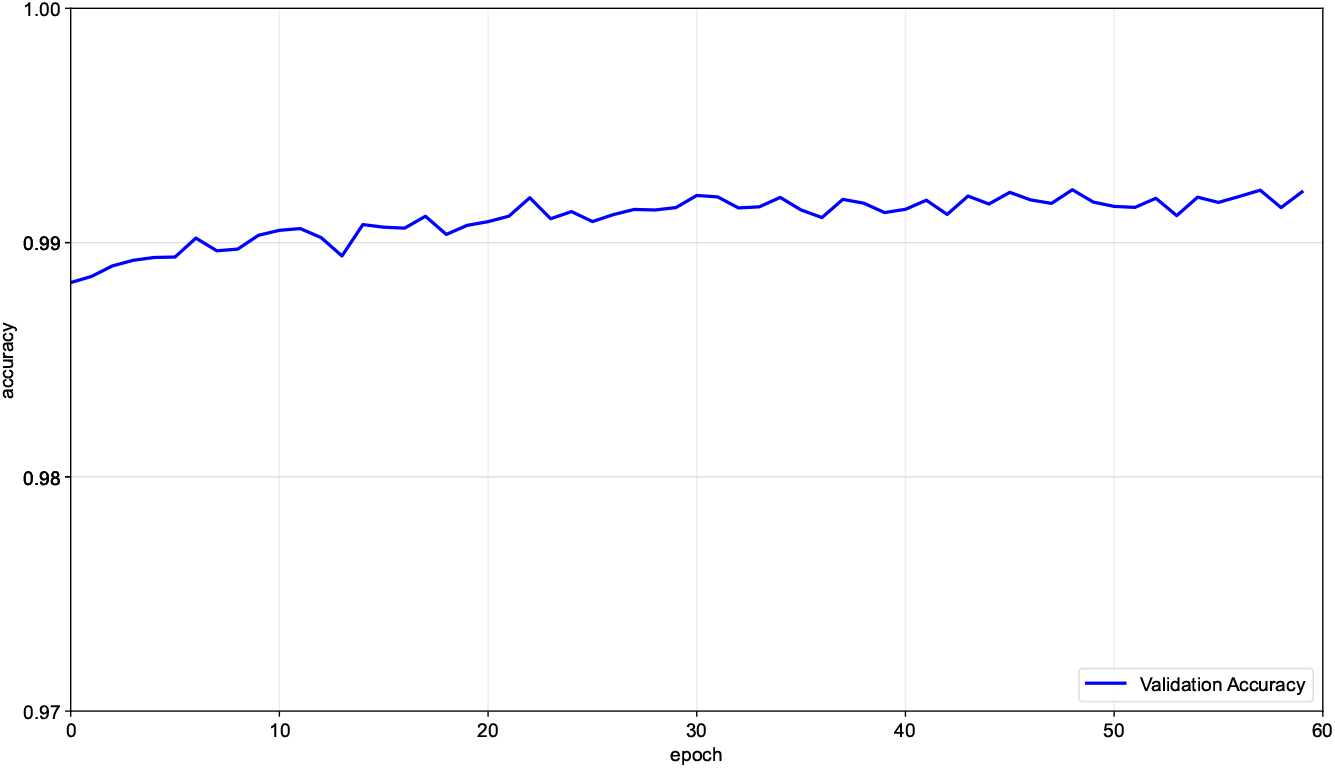
Validation Accuracy curve during training on the oversampled dataset.

The results demonstrate that the proposed model achieves highly competitive performance across both balanced and oversampled datasets. The oversampling strategy enhances recall and F1-score, improving detection of rare protein–protein interactions while maintaining strong precision. Consistently high AUC-ROC values further validate the model’s robust classification ability, and the confusion matrices in Fig 3 and Fig 4 show that most predictions fall along the true positive and true negative axes, indicating minimal misclassification. These findings establish that the model is effective on well-balanced data and remains reliable under imbalanced conditions, making it a promising framework for large-scale protein–protein interaction discovery and downstream bioinformatics applications.

**Fig 3.**
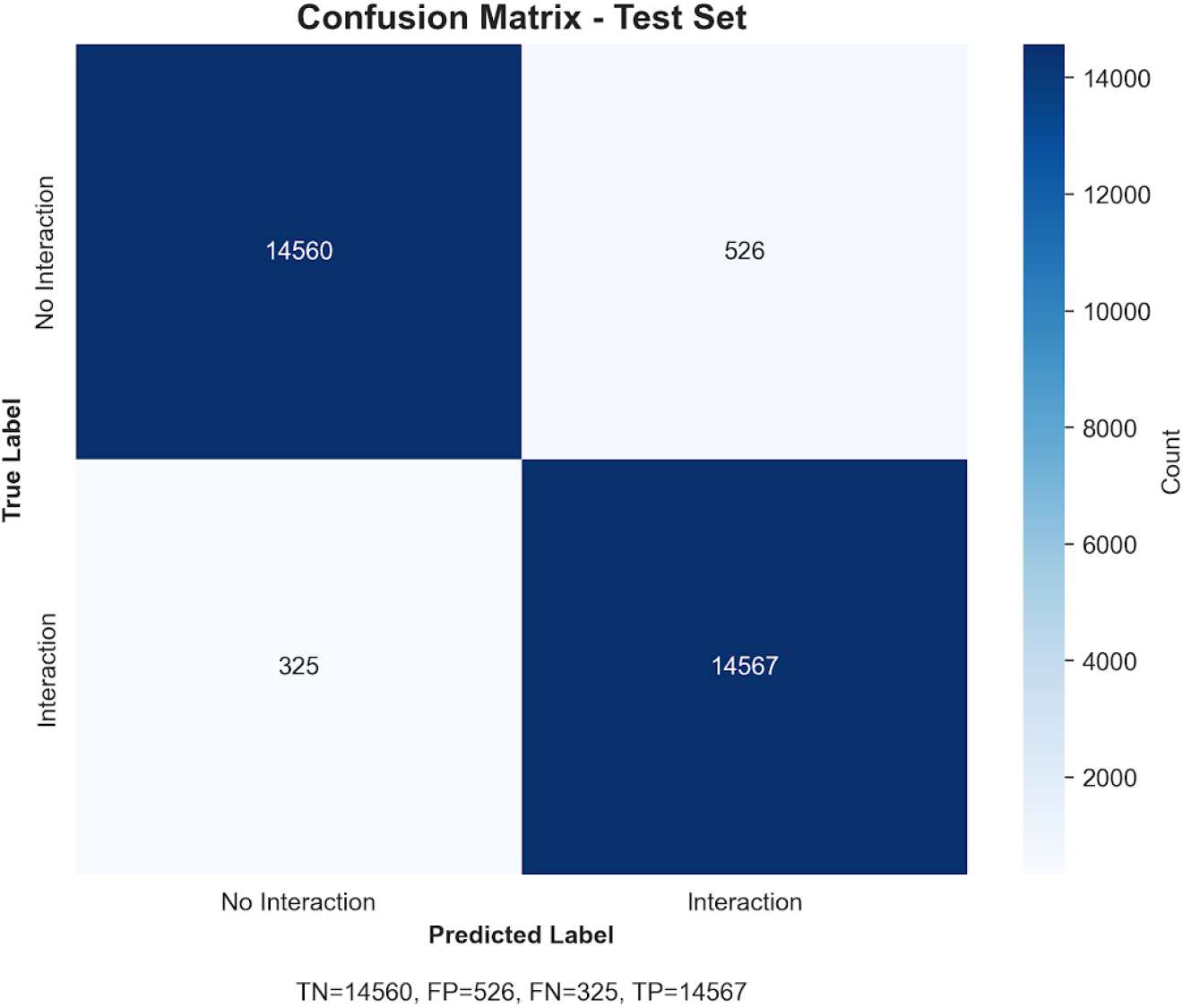
Confusion matrix on the balanced dataset.

**Fig 4.**
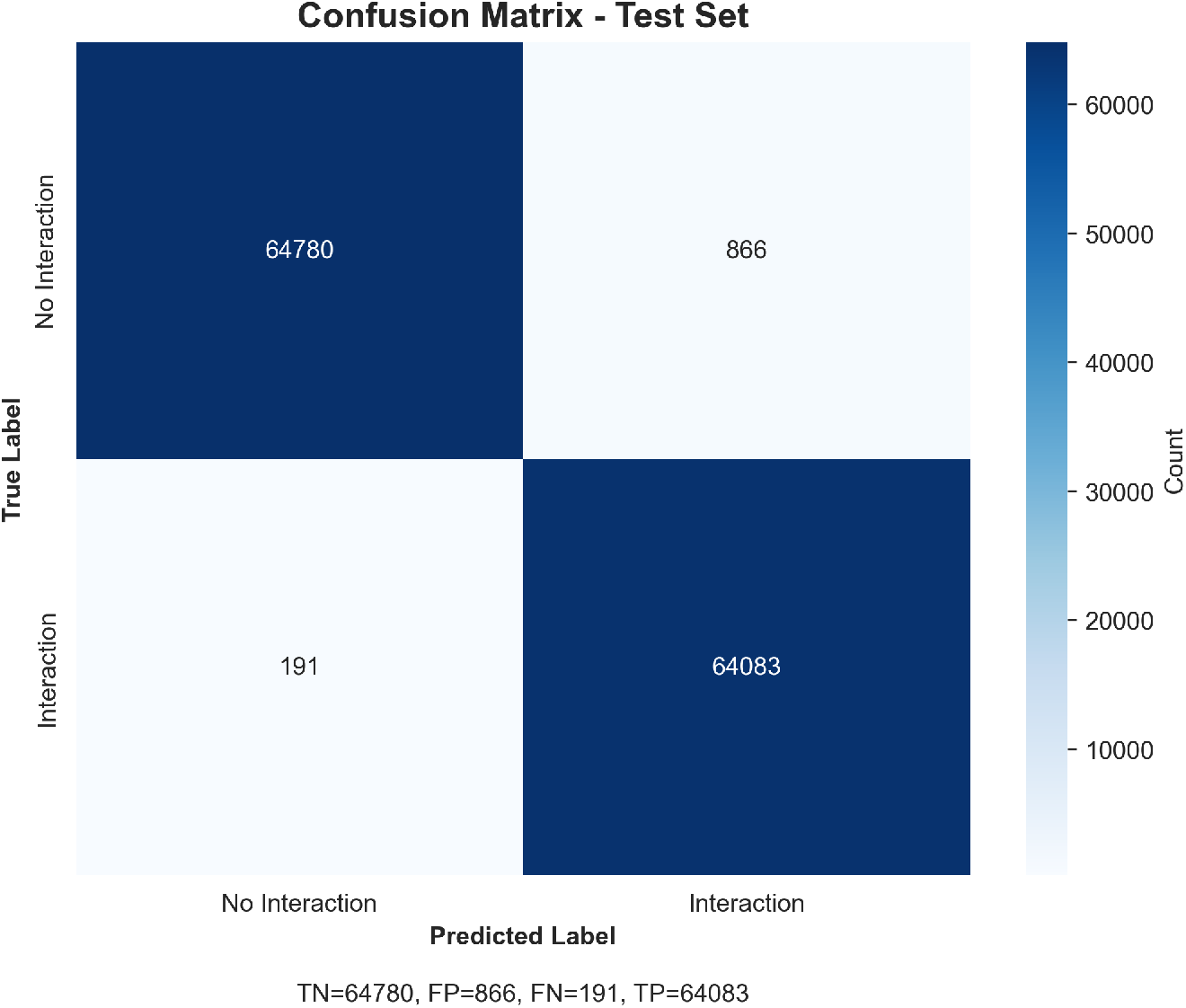
Confusion matrix on the oversampled dataset.

The performance of the proposed ProtAttn-QuadNet model was compared against several state-of-the-art protein-protein interaction prediction methods across multiple species and datasets (Table 2).

**Table 2.** Comparison of ProtAttn-QuadNet model with existing state-of-the-art models.

| Method | Dataset Details | ACC (%) | Precision (%) | Recall (%) | F1 (%) |
| --- | --- | --- | --- | --- | --- |
| DeepPPI (2017) [16] | *Saccharomyces cerevisiae* – 17,257 pos., 48,594 neg. | 92.50 | 94.38 | 90.56 | 92.52 |
| PIPR (2019) [18] | *S. cerevisiae* – 5,594 pos., 5,594 neg. | 97.09 | 97.00 | 97.17 | 97.09 |
| ADH-PPI (2022) [26] | *S. cerevisiae* – 5,594 pos., 5,594 neg. | 95.73 | 95.75 | 93.94 | 94.84 |
|  | *Helicobacter pylori* – 1,458 pos., 1,365 neg. | 92.63 | 92.84 | 96.09 | 94.44 |
| SemiGNN-PPI (2023) [27] | *Homo sapiens* (SHS27K) – 12,517 total | – | – | – | 92.40 |
|  | *H. sapiens* (SHS148K) – 44,488 total | – | – | – | 89.51 |
| xCAPT5 (2024) [22] | *H. pylori* – 1,458 pos., 1,365 neg. | 97.27 | 97.30 | 97.07 | 97.18 |
|  | *S. cerevisiae* – 5,594 pos., 5,594 neg. | 99.76 | 99.76 | 99.75 | 99.37 |
|  | *H. sapiens* – 27,593 pos., 34,298 neg. | 99.77 | 99.75 | 99.75 | 99.62 |
| SCMPPI (2024) [28] | *S. cerevisiae* – 11,188 total | 94.55 | 96.68 | 92.24 | 94.40 |
| HI-PPI (2025) [29] | *H. sapiens* (SHS27K) – 12,517 total | 85.82 | – | – | 79.57 |
|  | *H. sapiens* (SHS148K) – 44,488 total | 89.32 | – | – | 84.12 |
| ProtAttn-QuadNet (Ours) | Cross-Species (*UniProt*) – 541,331 pos., 541,331 neg. | 99.19 | 98.66 | 99.70 | 99.18 |

Across all evaluated datasets, ProtAttn-QuadNet demonstrated consistently superior predictive performance. For the large-scale cross-species UniProt dataset, which contains over 541,000 positive and 541,000 negative interactions, ProtAttn-QuadNet achieved an accuracy of 99.19%, precision of 98.66%, recall of 99.70%, and an F1-score of 99.18%, outperforming existing models. Notably, even on smaller organism-specific datasets, such as *S. cerevisiae* and *H. pylori*, ProtAttn-QuadNet either matched or exceeded the highest reported F1-scores from previous studies, including xCAPT5 and SCMPPI.

These results highlight the ability of ProtAttn-QuadNet to generalize across diverse PPI datasets while maintaining high precision and recall, which is critical for minimizing both false positives and false negatives in large-scale interactome predictions. Compared to earlier models, which often exhibited trade-offs between precision and recall depending on dataset size or species, ProtAttn-QuadNet achieves a balanced improvement across all evaluation metrics.

### Statistical Analysis

Comprehensive statistical analyses were performed to rigorously evaluate the reliability and significance of the proposed PPI prediction model. Three primary statistical tests were applied: the Chi-square test to assess statistical associations, the Wilcoxon signed-rank test to compare model performances, and an effect size analysis to determine the magnitude of observed differences. All statistical tests were conducted at a significance level of 0.05.

The Chi-square (*χ*^2^) test was applied to examine whether a statistically significant association existed between the predicted and true PPI classes (i.e., interacting vs. non-interacting protein pairs). The hypotheses were defined as:

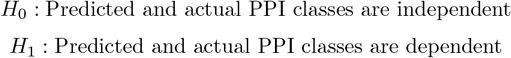

The Chi-square statistic was computed as:

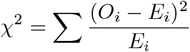

where *O*_*i*_ and *E*_*i*_ represent the observed and expected frequencies, respectively.

The analysis produced a *χ*^2^ value of 26,671.80 with 1 degree of freedom, *p <* 0.000001, and a Cramér’s *V* of 0.944, indicating a very strong statistical association between the predicted and actual PPI outcomes.

Fig 5 illustrates the percentage distribution across confusion matrix categories, clearly showing the deviation from random classification behavior.

**Fig 5.**
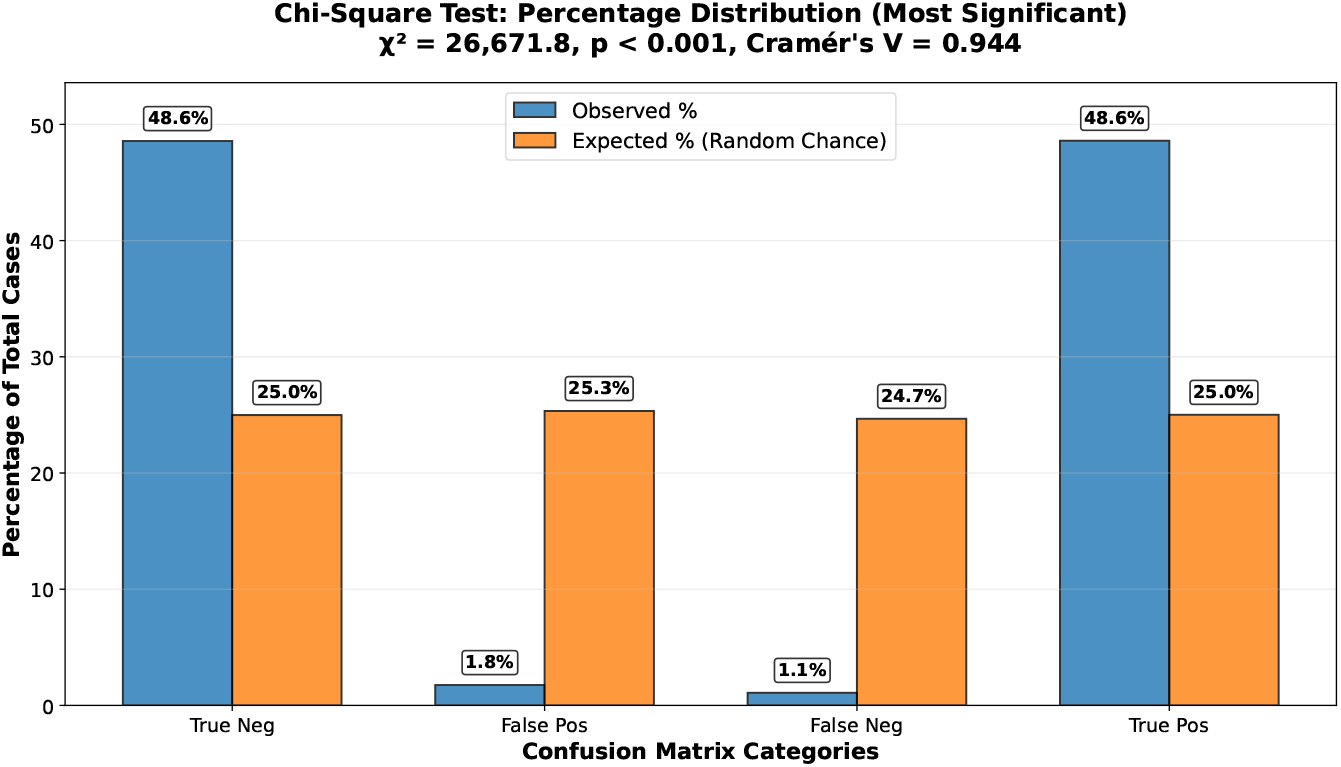
Distribution of predicted versus actual protein–protein interaction categories.

To confirm that the proposed PPI model significantly outperformed the random-chance baseline of 50%, we applied a Wilcoxon signed-rank test on the distributions of accuracy, F1-score, and AUC-ROC. Each metric yielded a test statistic of *W* = 990.0 with *p* = 5.68 *×* 10^*−*14^, indicating that the model’s performance gains were highly significant. Building on this, we quantified the strength of association and the magnitude of the predictive effect using multiple effect size measures. Cramér’s *V*,

Cohen’s *w*, and the Phi coefficient were all 0.944, reflecting a *very large* effect size. These results collectively demonstrate that the proposed PPI model not only significantly outperforms a random baseline but also produces predictions that are strongly consistent with true interaction labels, highlighting its practical and statistical relevance.

## Materials and Methods

### Dataset

A total of 573,661 protein entries were collected from the UniProtKB database [25]. Proteins were divided into interacting and non-interacting groups. Duplicate interaction pairs were removed; if both A=B and B=A were present, only one was kept. Each protein pair was labeled 1 for interaction (positive) or 0 for non-interaction (negative). To balance the dataset, two approaches were applied: (a) selecting an equal number of positive and negative samples, resulting in 249,814 protein pairs (124907 positive, 124907 negative) and 157,839 unique proteins, and (b) oversampling the positive class to include all proteins, resulting in 1,082,662 protein pairs (541331 positive, 541331 negative) and 573,661 unique proteins.

A separate dataset of 573,661 protein sequences was used for sequence embedding. Each protein sequence was converted into a 1024-dimensional numerical vector using ProtBERT embeddings [20], which were then used as model input.

### Data Preprocessing

In this study, robust scaling [30], defined in Eq. 5, was used to normalize protein embedding vectors. Biological data such as protein embeddings often exhibit non-Gaussian distributions, contain outliers, and represent diverse protein families, making standard normalization approaches less effective. Traditional z-score normalization, which relies on the mean and standard deviation, performs poorly when extreme values from rare proteins or functional domains distort statistical estimates. Protein embeddings derived from language models or structural encoders frequently show heavy-tailed distributions, with outliers originating from unusual structures, membrane domains, or disordered regions. Such outliers can negatively impact mean-based normalization, resulting in suboptimal model convergence and degraded performance.

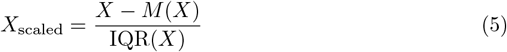

where *X* ∈ ℝ^*N ×D*^ is the protein embedding matrix with *N* proteins and *D* = 1024 dimensions, *M* (*X*) is the median of *X* across the protein dimension, and IQR(*X*) is the interquartile range of *X*.

### Dataset Splitting

A stratified splitting strategy was employed to ensure balanced class distribution across training, validation, and test sets while maintaining statistical rigor for model evaluation. To achieve this, a two-stage stratified sampling approach was adopted instead of a single-stage three-way split. Single-stage splitting can introduce subtle class imbalances due to rounding effects when dividing samples into three partitions simultaneously, whereas two-stage splitting manages one binary decision at a time (train+validation vs. test, then train vs. validation), ensuring precise stratification.

In the first stage, a fully isolated test set (12%) was created and kept untouched during model development, hyperparameter tuning, and validation to prevent data leakage. The second stage applied a test size of 0.225 to the remaining 88% of data, yielding a validation set that accounted for exactly 20% of the total dataset (0.225 *×* 0.88 ≈ 0.20). This method preserved nested stratification, maintaining the original class distributions at both stages—something not guaranteed by a single three-way split.

Formally, given a dataset *D* with binary interaction labels *Y* = *y*_1_, *y*_2_, …, *y*_*n*_, where *y*_*i*_ ∈ 0, 1, stratified splitting was performed as described above to preserve the proportions of positive and negative interactions across all partitions.

Stage 1: Training+Validation vs Test Split

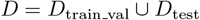

Stage 2: Training vs Validation Split

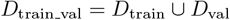

Thus, the final balanced dataset contains 169,873 training samples, 49,963 validation samples, and 29,978 test samples, while the oversampled dataset contains 736,210 training samples, 216,532 validation samples, and 129,920 test samples.

### Feature Engineering

Our objective is to enhance interaction-aware representations that capture diverse functional dependencies in protein–protein interactions (PPIs). We introduce two key feature types, Element-wise Product and Absolute Difference, which collectively capture complementary interaction dynamics.

1. **Element-wise Product** focuses on synergistic relationships by emphasizing dimensions where both proteins exhibit strong activations simultaneously. When both embeddings express high values in the same dimension, their product yields a large value, reflecting cooperative or co-regulatory behavior.
2. For protein embeddings **x**_1_, **x**_2_ *∈* R^*d*^, we define the interaction feature as:
3. 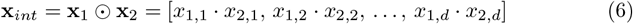
4. where, **x**_*int*_ captures *synergistic activation*, representing cooperative dimensions where both proteins contribute strongly.
5. **Absolute Difference** on the other hand, models complementary relationships by quantifying divergence between embeddings. Large values indicate distinct functional behaviors or differing biological characteristics between the two proteins.
6. We define a difference feature as:
7. 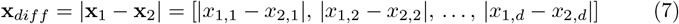
8. where, **x**_*diff*_ captures *complementary activation*, representing the magnitude of contrast across feature dimensions.
9. 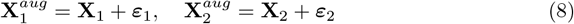

where **X**_1_, **X**_2_ ∈ ℝ^1024^ are the original scaled embeddings, ***ε***_1_, ***ε***_2_ *∼ 𝒩* (0, *σ*^2^*I*) are independent Gaussian noise vectors, *σ* = 0.02, and *I ∈* R^1024*×*1024^ is the identity matrix.

### Model Architecture

A multi-stream attention model is designed to predict protein–protein interactions while also estimating interaction uncertainty, binding strength, and interaction type. The model takes four different types of features as input and processes them through parallel attention streams and cross-attention layers to capture both individual protein properties and their relationships. The overall architecture of the proposed model is illustrated in Fig 6.

**Fig 6.**
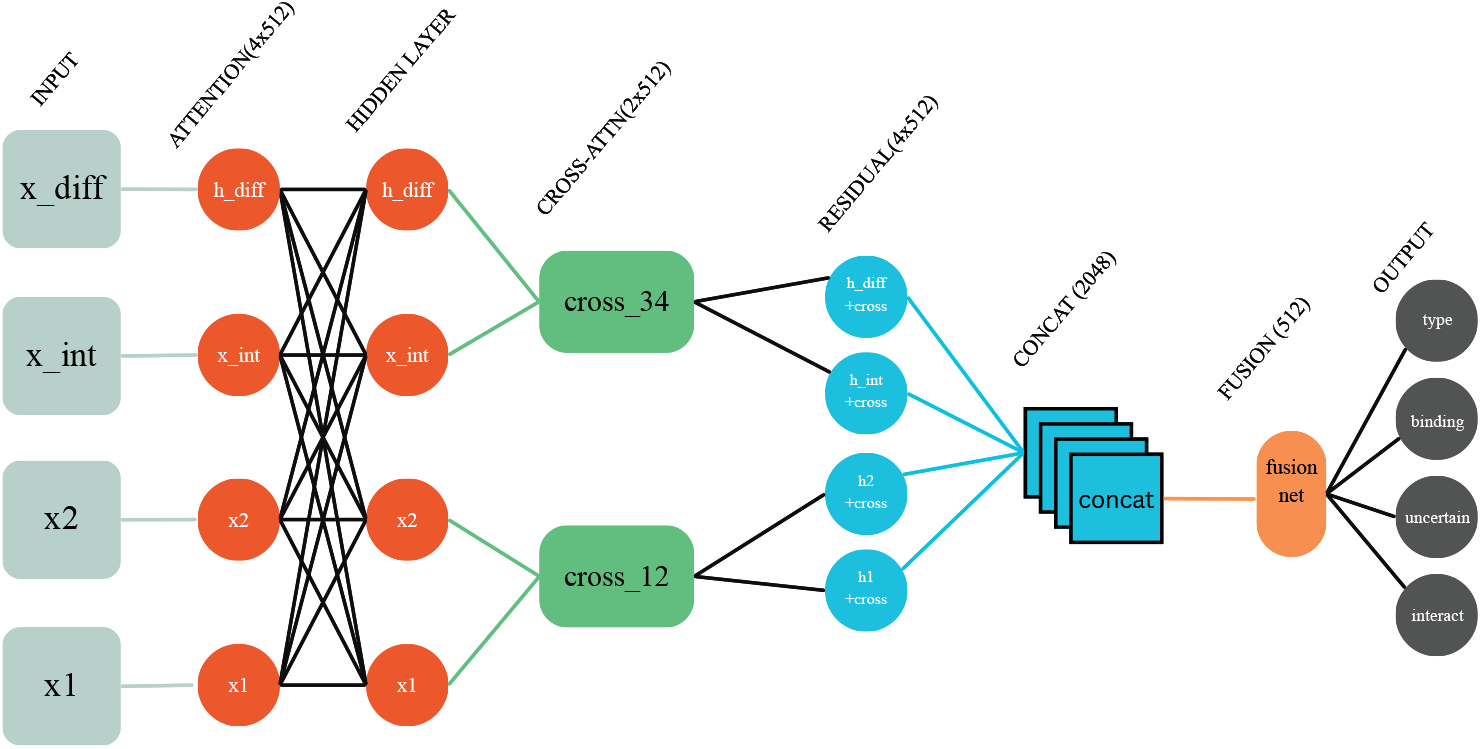
Architecture of the ProtAttn-QuadNet framework. Four distinct protein feature representations are processed through parallel attention streams and cross-attention layers to predict protein–protein interactions and simultaneously estimate interaction uncertainty, binding strength, and interaction type.

### Advanced Attention Block

The Advanced Attention Block is designed as the core computational unit of the architecture, which processes each feature stream through a sophisticated attention mechanism. Given an input **x** ∈ ℝ^batch size*×*input dim^, it is first projected into a higher-dimensional hidden space as follows:

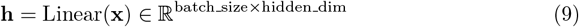

The projected features are normalized using layer normalization:

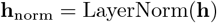

Subsequently, multi-head self-attention is applied:

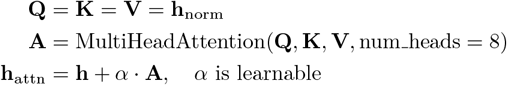

A feed-forward network with GELU activation is then applied:

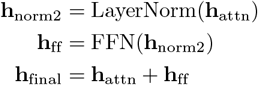

In this work, multi-head self-attention is employed to enable the model to focus on multiple aspects of each input dimension:

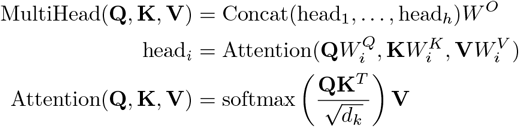

### Four-Stream Processing

The proposed architecture processes four feature types through parallel attention streams:

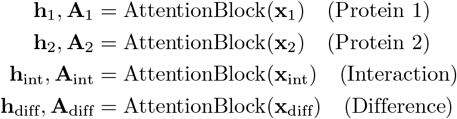

Each attention stream is allowed to focus on the most relevant features for its specific input type. This design facilitates specialized processing of individual protein properties, interactions, and differences, while maintaining a consistent architecture across all streams.

### Cross-Attention Fusion

To enable information exchange between streams, cross-attention mechanisms are implemented as follows:

Protein Cross-Attention:

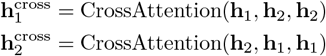

Feature Cross-Attention:

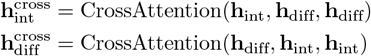

Residual connections are applied to the final stream representations to preserve the original information:

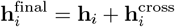

### Quad-Stream Fusion

After the final representations of each stream are computed, they are concatenated to form a combined feature matrix:

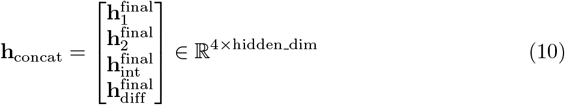

The concatenated representation is then passed through a two-layer fusion network with GELU activation, which reduces dimensionality while preserving critical information:

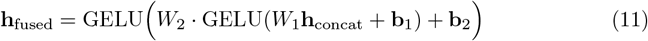

where the weight matrices and their dimensions are explicitly defined as:

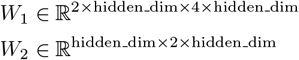

and **b**_1_, **b**_2_ are the corresponding bias vectors.

This fusion network integrates information from all four streams Protein 1, Protein 2, Interaction, and Difference into a single, compact representation while retaining the most important features for downstream prediction tasks.

### Multi-Task Learning Framework

A multi-task learning framework is employed to simultaneously predict multiple aspects of protein–protein interactions from the fused representation **h**_fused_. Predictions for four related tasks are generated from the fused representation:

1. **Interaction Prediction Head:** The probability of interaction is predicted using a sigmoid activation:

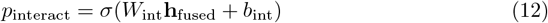
2. **Uncertainty Quantification Head:** The uncertainty of the interaction prediction is estimated using a Softplus activation:

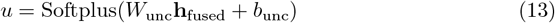
3. **Binding Strength Head:** The binding strength of the interaction is predicted using a sigmoid activation:

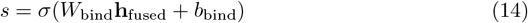
4. **Interaction Type Head:** The type of interaction is classified using a Softmax activation:

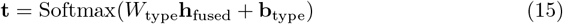

These task-specific heads are designed to enable the model to simultaneously learn complementary objectives, improving both predictive performance and representation learning across tasks.

### Loss Function Design

We define a multi-task loss that combines four components with carefully tuned weights:

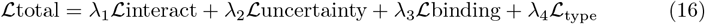

Here, ℒinteract represents the binary cross-entropy loss used to optimize the interaction prediction task. ℒuncertainty is a calibration-based term that penalizes overconfident or inconsistent predictions. ℒbinding employs mean squared error to estimate the binding strength between protein pairs. Finally, ℒtype models the interaction category using KL-divergence. The weights (*λ*_1_, *λ*_2_, *λ*_3_, *λ*_4_) are determined through hyperparameter tuning to balance the learning objectives effectively.

### Optimization and Hyperparameter Search Strategy

A systematic grid search integrated with advanced optimization techniques is employed to determine the optimal configuration of the ProtAttn-QuadNet architecture. The search space includes optimizer selection, learning rate, weight decay, batch size, learning rate scheduling, and multi-task loss weighting.

Four learning rates (0.0001, 0.0005, 0.001, 0.002), three weight decays (1 × 10^*−*5^, 1 × 10^*−*4^, 1 × 10^*−*3^), three optimizers (Adam, AdamW, RMSprop), three schedulers (ReduceLROnPlateau, CosineAnnealing, StepLR), and four batch sizes (32, 64, 128, 256) are explored. Additionally, three pre-configured settings are evaluated to assess stability and performance consistency.

- Conservative: LR=0.0005, WD=1e-4, AdamW, ReduceLROnPlateau
- Aggressive: LR=0.001, WD=1e-3, Adam, CosineAnnealing
- Balanced: LR=0.0001, WD=1e-5, RMSprop, StepLR

The best performing configuration is presented in Table 3.

**Table 3.** Optimal Configurations for Balanced and Oversampled Datasets.

| Parameter | Balanced Dataset | Oversampled Dataset |
| --- | --- | --- |
| Optimizer | AdamW | RMSprop |
| Learning Rate | 0.0005 | 0.0001 |
| Weight Decay | $1 \times 10^{-4}$ | $1 \times 10^{-5}$ |
| Scheduler | ReduceLROnPlateau | StepLR |
| Batch Size | 128 | 256 |

## Data availability

The primary data is available at https://www.uniprot.org/. The processed data and code will be archived on GitHub.

